# Human ADME/PK is lost in translation and prediction from *in silico* to *in vitro* to *in vivo*

**DOI:** 10.1101/2025.02.17.638712

**Authors:** Urban Fagerholm, Sven Hellberg

**Affiliations:** Prosilico AB, Lännavägen 7, SE-141 45 Huddinge, Sweden

## Abstract

**Background:** Measurements and predictions of aqueous solubility (S), apparent cell permeability (P_app_), unbound microsomal intrinsic clearance (CL_int,u_), unbound fraction in plasma (f_u_), log D, Lipinski’s Rule of 5 (Ro5) and BBB+/BBB- (blood-brain-barrier) are commonly used in early drug discovery to evaluate whether compounds are likely to have adequate ADME/PK in humans. The main objective was to evaluate the validity and proposed thresholds for commonly used *in silico* to *in vitro* (with particular interest in the ADMET Predictor software) and *in vitro* to *in vivo* human ADME/PK prediction methods, and physicochemical estimates and rules of thumb. A secondary aim was to compare validity and thresholds to that for the prediction software ANDROMEDA.

**Methods:** Data were collected from literature and own studies. Main measures of validity were Q^2^ (true predictive accuracy; *in silico* models), R^2^ (correlation coefficient; *in vitro* models), Q^2^ • R^2^ (for translation from *in silico* to *in vivo* via *in vitro*), skewness, range, % correct predicted class and clinical relevance.

**Results and Discussion:** Poor accuracies (Q^2^ • R^2^=0.05, 0.05, 0.36 and 0.45; ∼0 at low to moderate, decision-critical levels) and class predictions, limited ranges (not covering low to moderate estimates), systematic errors (often considerable overprediction at low values), poor clinical relevance and inadequate thresholds were found. Predictive accuracy was mainly lost in the translation from *in vitro* to *in vivo*. Log D and Ro5 are poor predictors of oral bioavailability and half-life. Ro5 produced 63 % false negatives for prediction of poor/good oral absorption. The overall mean Q^2^ for ANDROMEDA was 3 times higher (0.57 *vs* 0.2). Advantages with ANDROMEDA compared to *in vitro* data-based *in silico* models include wider application domain (high resolution at low values), extrahepatic elimination models, minimal skewness, clinical relevance, balanced thresholds, and compound- and parameter-unique confidence intervals. ANDROMEDA successfully predicted the human clinical ADME/PK for all small drugs marketed in 2021, while the *in silico* to *in vitro* to *in vivo* approach was out of reach for all.

**Conclusion:** The validity of investigated methodologies (including ADMET Predictor) and thresholds were overall very low. Unless both predicted S, permeability and f_u_ are high and CL_int_ is moderate (an overall criterium not met for investigated modern small drugs) one is more or less lost in the translation, and will jeopardize compound selection and optimization. This clearly shows the need for better and thoroughly validated prediction models and software. Marked improvements in accuracy, range, balance and clinical relevance were achieved with ANDROMEDA, which predicts human clinical ADME/PK directly from chemical structures and has undergone extensive validation.

## INTRODUCTION

*In vitro* data, physicochemical parameter estimates and rules of thumb, and classification systems, are commonly used in early drug discovery to predict and evaluate whether compounds are likely to have adequate or unfavorable ADME/PK properties in humans.

Examples of *in vitro* and physicochemical parameters include aqueous solubility (S), apparent Caco-2 and MDCK permeability (P_app_), unbound microsomal intrinsic clearance (CL_int,u_), unbound fraction in plasma (f_u_), log P and D (typically at pH 7.4), molecular weight (MW), and number of hydrogen bonding acceptors (HBA) and donors (HBD). Commercially available software for ADME/PK predictions (mainly *in silico* to *in vitro*) include ADMET Predictor by Simulations Plus Inc. and QikProp by Schrödinger.

Physicochemical rules of thumb include proposed ideal (for ADME/PK) log D and log P of 0-3 (or 2-3) and 1-5 [1–3], respectively, S>10-200 µM for acceptable *in vivo* solubility/dissolution [4, 5], low microsomal CL_int,u_ (high metabolic stability; <ca 10-15 µL/min/mg) [6, 7], and maximum 1 violation of criteria of Lipinski’s Rule of 5 (Ro5, violation criteria are MW>500 g/mol, log P>5, >10 HBA and >5 HBD) for acceptable gastrointestinal uptake and oral bioavailability [8].

Classification systems include BCS (Biopharmaceutics Classification System; 4 classes depending on high and low S and permeability) and BBB+/BBB- (BBB=blood-brain barrier). In the BCS, compounds in Classes II and III have, or are predicted to have, solubility/dissolution rate-limited and permeability-limited gastrointestinal uptake in humans *in vivo*, respectively, whereas compounds of Class I possess, or are predicted to have, good/sufficient solubility/dissolution and permeability, and Class IV-compounds have, or are predicted to have, gastrointestinal uptake limited by both solubility/dissolution and permeability [9]. BCS-classing is commonly done based on measured S at various pH-levels) and P_app_-estimates, and criteria/thresholds for these.

According to BBB+/BBB-, compounds belonging to BBB+ (proposed to have high log equilibrium brain/blood concentration ratio (log BB; various thresholds suggested)) are believed to easily cross the BBB and enter the brain rapidly (sufficient brain uptake and distribution is essential for centrally acting compounds), whereas BBB- compounds are anticipated/predicted to not cross the BBB or slowly and poorly cross the BBB and enter the brain (favorable if no central side effects are wanted).

*In silico* prediction models have been developed for, for example, log D, log S, log P_app_, log human effective permeability (P_eff_), fraction absorbed *in vivo* (f_abs_), microsomal log CL_int,u_, lin and log f_u_ and log BB (BBB+/BBB-) [10–17], which enables ADME/PK predictions for compound-selection and -optimization, and decision making even before compound synthesis.

Sufficient validity (good accuracy, no or minor systematic trends, wide range and high clinical relevance), reliability (high reproducibility), and adequate proposed thresholds are required for *in silico* and *in vitro* prediction methods and adequate compound selection. To our knowledge, an evaluation of this for commonly used *in silico* and *in vitro* methods, physicochemical parameters (including proposed rules of thumb) and classifications has not been done before. Based on the costs and time spent and failures in drug discovery and development, and wish/requirement to replace, reduce and refine animal experiments (3R) and to assure safety in clinical trials, this is surprising. Lack of concern by users, widespread perception that it is an area with sufficiently accurate methodology, unwillingness by model and software developers to disclose poor results and clinical relevance, and lack of interest and demands from regulatory agencies, are among possible explanations.

Another approach (than *in vitro* data-based *in silico* prediction models and physicochemical parameter estimates) is *in silico* ADME/PK-models based on human clinical parameters and estimates. Our team (Prosilico AB, Sweden) has developed and validated such a prediction, simulation and optimization software, ANDROMEDA (https://prosilico.com/andromeda). ANDROMEDA is built based on a unique, large data bank, unique algorithms, and conformal prediction and machine learning methods. It produces valid levels of confidence and reaches an average Q^2^ (correlation between measured and predicted estimates in cross-validations) of 0.52 for its numerical models and parameters within the major domain of MW 150-750 g/mol. Its *in silico* models for permeability-based fraction absorbed (f_abs,pe_; range 0-100 %), fraction dissolved *in vivo* (f_diss_; range 1-100 %), log CL_int_ (range 1-200,000 mL/min) *in vivo* in man, and *in vitro* log f_u_ (f_u_-range 0.02-100 %), have minimal skewness and Q^2^-values of 0.7-0.8, 0.55, 0.55 and 0.6-0.7 (0.5-0.55 for lin-lin scale), respectively [18–21]. How well other established *in silico*-to-*in vitro*- to-*in vivo* prediction approaches (as above) perform compared to ANDROMEDA (directly from *in silico* to *in vivo* in man, without using *in vitro* data) has not been fully investigated and shown.

The main objective of this study was to evaluate the validity and proposed thresholds for commonly used *in silico* to *in vitro* and *in vitro* to *in vivo* methods, and physicochemical estimates and rules of thumb, for prediction of human clinical ADME/PK. Results obtained with the commercially available software ADMET Predictor by Simulations Plus Inc. were of particular interest. A secondary aim was to compare the validity and thresholds to that for ANDROMEDA.

## METHODS

### Sources of Data

The following references were used: log D [10, Prosilico data on file], log S and f_diss_ [10, 13, 14, 22-25, Prosilico data on file], log Caco-2 P_app_ [10, 15, 16, 23, 25, 26–30], log MDCK P_app_ [31], log P_eff_ (ADMET Predictor predicts P_eff_ and not P_app_) [10, 31, 32], log microsomal CL_int,u_ [6, 7, 10, 30, 31-35, Prosilico data on file], f_u_ (13, 20), Ro5 [36], log BB and BBB+/BBB- [37–39], and *in vivo* f_abs_ [40]. *In silico* to *in vitro* ADME/PK results obtained using ADMET Predictor by Simulations Plus Inc. were taken from references [10], [14] and [17].

### Data Analysis

Measures of validity were Q^2^ (accuracy; correlation between measured and predicted estimates in cross-validations, for test or hold-out sets), mean absolute error (MAE), root mean square error (RMSE), skewness (Q^2^ and intercepts on axis for observed/measured values), range (minimum to maximum observed and predicted values), % correct predicted class and clinical relevance (including analysis of similarities and differences between *in vitro* and *in vivo* parameters and conditions, as well as levels of accuracy).

The predictive performance of *in vitro* methods is often demonstrated as R^2^-values (correlation coefficient for measured *vs* predicted estimates; retrospective/non-prospective), where values <limit of quantification (LOQ) have been excluded.

Proposed thresholds and optimal ranges and limits of quantification for *in vitro* and physicochemical parameters were compared to corresponding apparent *in vivo* values.

The translation from *in silico* to humans *in vivo* (Q^2^), via *in vitro* (R^2^), was done by multiplying the both estimates (Q^2^ • R^2^) for each parameter (log S to *in vivo* f_diss_, log P_eff_, via log P_app_, to *in vivo* f_abs_, and *in vitro* to *in vivo* CL_int_).

Results were compared to those obtained with ANDROMEDA for corresponding parameters.

## RESULTS

### In silico to in vitro Predictions

#### Log D

For a test set of 287 compounds with log D ranging from −3 to 4 (no data in the interesting and challenging region log D>4), ADMET Predictor produced Q^2^, MAE and RMSE for *in silico vs in vitro* log D of 0.79, 0.59 and 0.76, respectively [10]. Ca 3 % of compounds had 1.5- to 2-fold prediction errors.

#### Solubility (S)

The Q^2^, MAE and RMSE for *in silico vs in vitro* log S with ADMET Predictor (for a test set of 691 compounds; no or few with log D>4) were 0.20, 1.06 and 1.23, respectively [10].

Using a 4-class system (low/poor, moderate, high, very high), S-classification was correctly predicted for 56 % of compounds. Ca 95 % of low S compounds according to measurements (can be interpreted as undesirable/unacceptable S; <10 µM) were predicted to have moderate to very high S (desirable/acceptable S; >90 µM), whereas only ca 5 % of compounds with very high S were predicted to have low to high S (Figure 1). Ca 8 % of compounds with predicted very high S (desirable/acceptable S) had low measured S (undesirable/unacceptable S) and every other compound predicted low S did not have low measured S. Ca 67 % of compounds with very high predicted S also had very high measured S.

**Figure 1.**
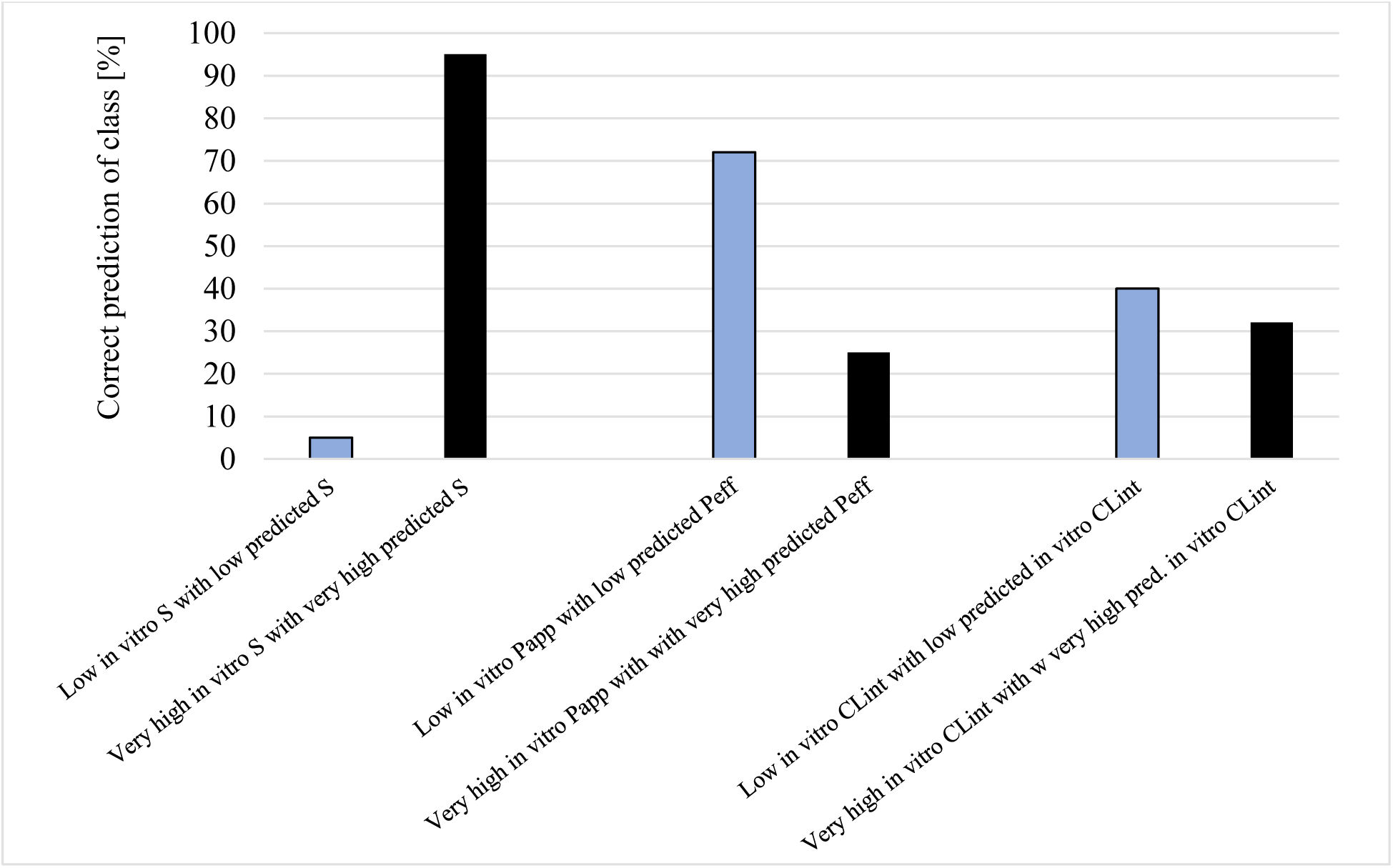
Percentage correct *in silico* predicted class (low and very high) for *in vitro* S, P_app_ & P_eff_ and CL_int,u_ for ADMET Predictor) [10].

Chevillard et al. [14] also used ADMET Predictor, and reached a Q^2^ of 0.32 and a RMSE of 1.25 for a test set from the *Solubility Challenge* (n=26; lowest S ca 0.1 µM). They also tested 10 other methods/software, including QikProp, and found Q^2^- and RMSE-values of 0.18-0.57 (0.34 for QikProp) and 1.09-1.66 (1.24 for QikProp) [13]. Corresponding results for another dataset (n=62) were 0.13-0.62 (0.56 for ADMET Predictor and 0.45 for QikProp) and 0.51-0.95 (0.72 for ADMET Predictor and 0.86 for QikProp). In a study by Falcón-Cano et al., Q^2^ of 0.4-0.8 and MAE of 0.6-0.8 were reached [14].

The Q^2^ for *in silico* to *in vitro* log S was estimated to average ca 0.4 (Figure 2).

**Figure 2.**
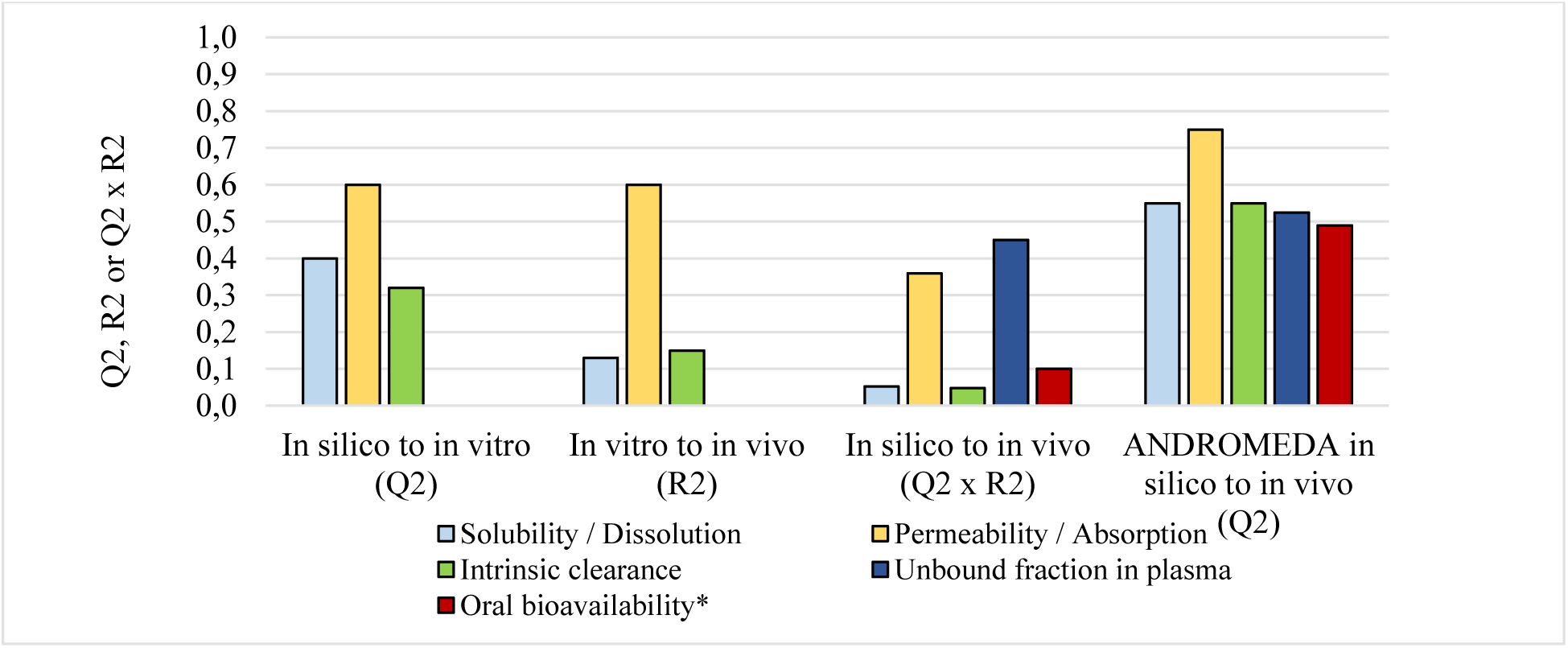
Q^2^ (true predictive accuracy; *in silico* models), R^2^ (correlation coefficient; *in vitro* models) and Q^2^ • R^2^ (for translation from *in silico* to *in vivo* via *in vitro*) for S to *in vivo* f_diss_, log P_eff_ to *in vivo* f_abs_ and microsomal CL_int,u_ to *in vivo* CL_int_, Q^2^ for *in silico* prediction of f_u_ and F (both ADMET Predictor), and predicted log D *vs* F. *Values for log D and ADMET Predictor were similar (ca 0.1) and combined. *Note: LOQ is a limitation for in vitro S, microsomal CL_int,u_ and f_u_, and therefore, many compounds with low to moderate estimates are missing and in silico prediction models for these show overprediction trends in these regions. Q^2^, R^2^ and Q^2^ • R^2^ for in vitro based models are much lower at low values*.

#### Apparent Caco-2 permeability (P_app_) and effective human permeability (P_eff_)

The % correct permeability classing and Q^2^ for *in silico* P_eff_ *vs in vitro* log P_app_ were 61 % and 0.61 (n=516) for a local model built by [10], and 47 % and unknown (roughly estimated to 40-50% based on the difference in correctly predicted permeability class) for ADMET Predictor [10], respectively. For a smaller set of commercially available compounds (n=26) the Q^2^ was 0.78 [10]. In cannot be ruled out that this set, or parts of it, was included in the training set for the model, and that these results, therefore, were exaggerated. Winiwarter et al. also reached Q^2^=0.60 [16]. Q^2^-values from 7 other studies were 0.16-0.74 [15]. The overall Q^2^ from the 10 studies was ca 0.6 (Figure 2). The impact of test compounds also present in training sets is unknown. The RMSE was estimated to 0.6-0.8, which corresponds to ca ±15-20 %, ±20 % and ±10 % variability of *in vivo* f_abs_ at P_app_ of ca 0.1, 1 and 10 • 10^-6^ cm/s (*in vivo* f_abs_ ca 30, 60 and 90 %, respectively [23]) [15]). At P_app_ corresponding to incomplete *in vivo* f_abs_ (<80-90 % f_abs_), there was virtually no correlation between measured and *in silico* predicted P_app_ (R^2^<0.1) [15]. For example, there were compounds with *in vitro* and *in silico* predicted f_abs_ of ca 80-90 % *vs* ca 10-15 % (7-fold difference) and ca 5-10 % *vs* 60 % (8-fold difference), respectively [15].

In 53 % of cases, incorrect *in vitro* P_app_-class was predicted using ADMET Predictor [10]. Ca 30 % of low/poor P_eff_ compounds (<1 • 10^-4^ cm/s; considered undesirable/ unacceptable permeability) were predicted to have moderate to very high P_app_ (>2 • 10^-6^ cm/s; desirable/acceptable permeability), and none of the compounds with predicted very high P_eff_ (>3 • 10^-4^ cm/s; desirable/acceptable permeability) had low measured P_app_ (<2 • 10^-6^ cm/s; undesirable/unacceptable permeability) (Figure 1). Ca 70 % of compounds with predicted low P_eff_ had low measured P_app_, ca 13 % of compounds with predicted low P_eff_ had very high measured P_app_ (>10 • 10^-6^ cm/s), and ca 67 % of compounds with predicted very high P_eff_ also had very high measured P_app_.

#### Human microsomal unbound intrinsic clearance (CL_int,u_)

ADMET Predictor produced a Q^2^ of 0.50 for *in silico vs in vitro* log CL_int,u_ for a test set consisting of 1199 compounds [10]. 37 % of predicted classifications were correct. Ca 68 % of very high CL_int,u_ compounds (undesirable/unacceptable CL_int,u_; >80 µL/min/mg) were predicted to have low to high CL_int,u_ (desirable/acceptable CL_int,u_), ca 18 % of compounds with predicted low CL_int,u_ (desirable/acceptable CL_int,u_; <10 µL/min/mg) had very high measured CL_int,u_ (undesirable/ unacceptable CL_int,u_), and ca 26 % of compounds with predicted very high CL_int,u_ (undesirable/unacceptable CL_int,u_) had low to high measured CL_int,u_ (desirable/acceptable CL_int,u_) (Figure 1). In studies by Shah et al. [6] (2 classes; low (ca <40-50 µL/min/mg) and high microsomal CL_int,u_) and Podlewska and Kafel [7] (3 classes; low (ca <10-15 µL/min/mg), medium and high microsomal CL_int,u_ (ca >40-60 µL/min/mg)), 80 and 70 % correct class predictions were achieved, respectively.

ADMET Predictor was also used to predict *in vitro* microsomal log CL_int,u_ for a set of 75 commercial reference compounds, and a Q^2^ of 0.14 was reached [10]. The average Q^2^ for ADMET Predictor for the two test sets was 0.32 (Figure 2).

#### Unbound fraction in plasma (f_u_)

Yun et al. used 3 different *in silico* methods, ADMET Predictor and models by Watanabe et al. and Ingle et al., to predict the *in vitro* f_u_ of 818 compounds with f_u_ ranging from 1 to 100 % (those with f_u_<1 %, which represent just over 10 % of all compounds and are extra challenging to measure and predict, were excluded) [17]. For models by Watanabe et al and Ingle et al, no test set compounds were used in the model training sets. This was, however, not assured for ADMET Predictor (implies risk of exaggerated outcome).

Q^2^-values (for linear predicted *vs* measured f_u_) for ADMET Predictor and models by Watanabe et al. and Ingle et al. were 0.52, 0.46 and 0.37, respectively [17]. Corresponding MAE- values were 13, 14 and 16 %, respectively. Clear trends were found. There was general overprediction at low (<25 %) f_u_ and underprediction at high f_u_. At f_u_ of 1-5 %, the average overprediction was ca 5-10 actual f_u_-% (ca >1 to 50-fold individual underpredictions), and at 90- 100 % f_u_ there was on average ca 2-fold underprediction.

### In vitro to in vivo Predictions

#### Solubility (S) to in vivo dissolution (f_diss_)

An investigation of the relationships between log S and *in vivo* f_diss_ and f_abs_ showed that these two measurements correlate poorly: R^2^=0.13 for log S *vs in vivo* f_diss_ (n=82) and R^2^=0.00 for log S *vs in vivo* f_abs_ (n=452) (Figure 3), respectively [22, 25]). The R^2^-estimates for S and dose-corrected log S *vs in vivo* f_abs_ were also zero.

**Figure 3.**
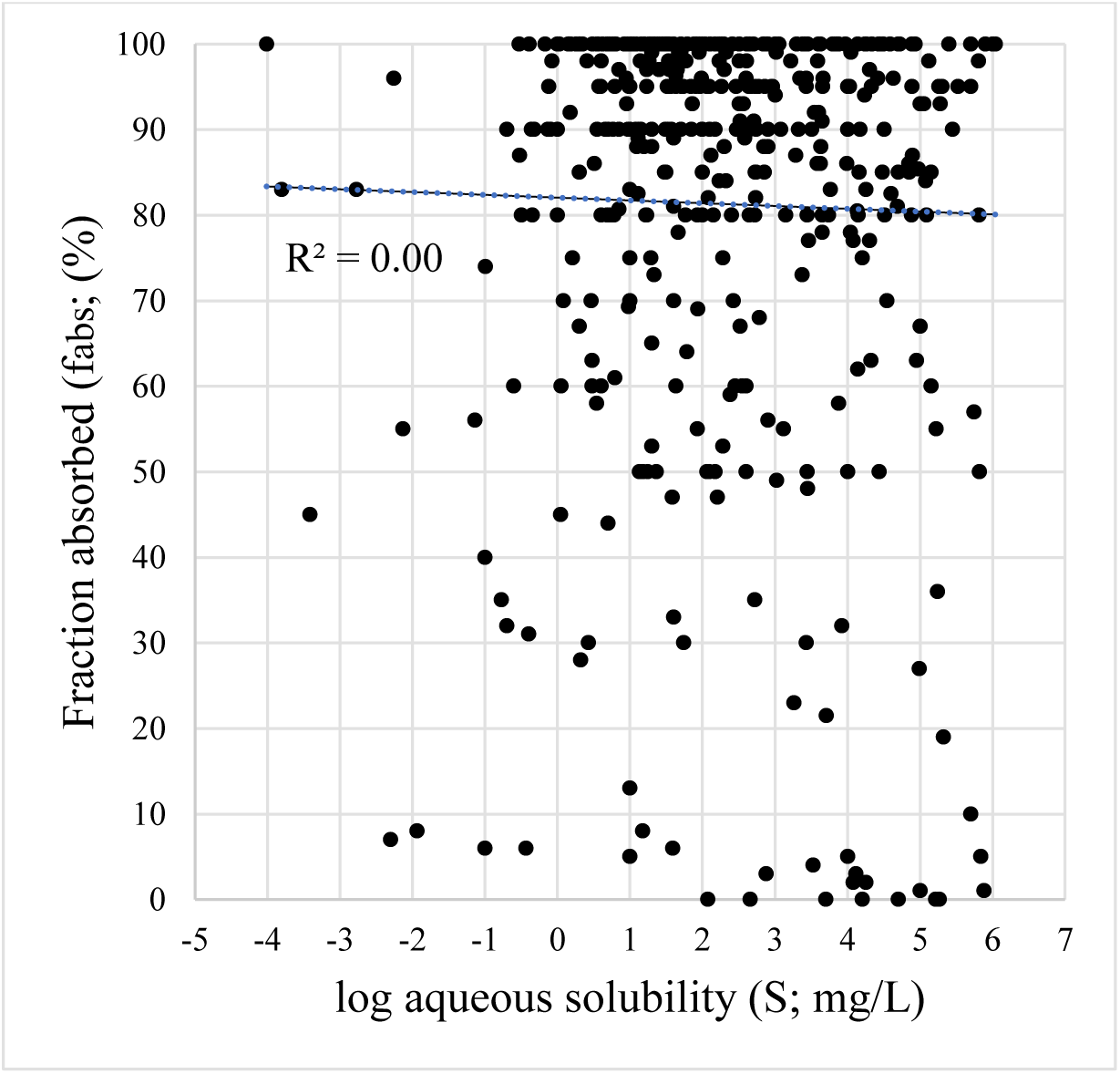
The correlation between measured log aqueous solubility (S) and fraction absorbed *in vivo* (f_abs_) for 452 compounds with log S ranging from −4.0 to 6.0 and f_abs_ ranging from ca 0 to 100 %.

For 70 compounds with S<5 mg/L (ca <10-20 µM) and 17 compounds with S<1 mg/L (ca <2-4 µM) there were weak correlations between log S and *in vivo* f_abs_ (R^2^=0.07 and 0.02, respectively) [25]. Complete or near complete f_abs_ has been demonstrated for many compounds with S∼2-4 µM, and moderate uptake has been shown for compounds with S∼0.2-0.4 µM [23, 25]. Out of 129 compounds of the *Solubility Challenge* data set [24], 27 have S in the 1-100 µM- range, and only one of these have an *in vivo* f_abs_ clearly limited/determined by solubility/dissolution [25]. The level of S corresponding to general solubility/dissolution-limited uptake *in vivo* in man (overall median <1 µM; individual <1-100 µM; approximated average S at f_diss_=80 % [25]) is low compared to proposed thresholds proposed (<10-200 µM [4, 5]; <10 µM in ADMET Predictor study by [10]).

The interlaboratory variability for S is large (>10- and >2-fold average and median ratio between maximum and minimum reported estimates, respectively) [25]. For example, there are 8400- and 4500-fold differences for highest and lowest reported S for dipyramidole and diclofenac (low to very high S for both), respectively [23].

For some low solubility drugs (at least 30 found), S has not been possible to quantify [41], including the well absorbed (high f_abs_) amlodipine, aripiprazole, bicalutamide, conivaptan, lomustine, posaconazole, venlafaxine and zolpidem, and poor f_diss_-compounds atavaquone, artemether, lumefantrine and paricalcitol. Thus, it has not been possible to predict their f_abs_ using S-data and establish *in vitro-in vivo* relationships along the whole S-scale. Almost every other compound with a non-quantifiable S, *in vivo* dissolution is or is predicted to be complete or near complete, and there at least over a handful of drugs with S>90 µM (very high *in vitro* S) are known to have incomplete *in vivo* dissolution.

In an investigation, 77, 84 and 100 % of compounds (n=73) predicted to be in BCS classes I (high P_app_ - high S), III (low P_app_ - high S) and IV (low P_app_ - low S) based on *in vitro* data also belonged to these classes based on their measured *in vivo* f_abs_, respectively [25]. For compounds that were predicted to belong to BCS class II (high P_app_ - low S), however, only 31 % belonged to *in vivo* BCS class II (69 % had an *in vivo* f_a_ of 90-100 %; BCS class I). For compounds that have BCS-classification from different sources (n=140), 64 % show contradictory classifications.

#### Permeability to fraction absorbed (f_abs_)

The R^2^ between log Caco-2 P_app_ and *in vivo* f_abs_ ranges between ca 0.2 and 0.8 (average ca 0.6; Figure 2), depending on compound choice, f_abs_-range and laboratory [26–30]. Limitations with *in vitro* permeability assays include greater uncertainty and quantification problems for low permeability compounds, low recovery (for example, for highly lipophilic compounds) and variability among labs. Skolnik et al. [26] found that 1/8 and 50 % of compounds were subject to <30 % (44 % of poorly soluble compounds) and <80 % recovery in Caco-2 experiments, respectively. In the literature, we have found >100 compounds that needed to excluded from evaluation of log Caco-2 P_app_ *vs in vivo* f_abs_-relationships for such reasons [41]. According to an evaluation and summary by Pham-Tee et al. [23], Caco-2 values corresponding to low f_abs_ (<30- 40 %) are ≤0.1 • 10^-6^ cm/s, and this is considerably lower than the threshold for low P_app_ (<2 • 10^-6^ cm/s; corresponding f_abs_ ca <65 %) set by Sohlenius-Sternbeck and Terelius who evaluated ADMET Predictor [10].

The method for estimation of *in vivo* P_eff_ has difficulties to accurately quantify for compounds with f_abs_<ca 70-90 % (similar as the f_abs_ of the average drug) and P_eff_ ca <1 • 10^-4^ cm/s [31, 32], which limits it for applicability to predict complete (ca 90-100 %) or incomplete uptake (ca <70-90 %).

In an evaluation of MDCK cell P_app_-based predictions of P_eff_ and human *in vivo* f_abs_ (similar to the ADMET Predictor approach) (n=46; all except one with apparent permeability limited uptake), a R^2^ of 0.43 and an intercept of 39 % f_abs_ were reached, respectively [31]. A considerable portion of the compounds (nearly 40 %) of test compounds were, however, already used in the training set, and therefore, the accuracy was most likely exaggerated.

#### Intrinsic clearance (CL_int_)

Our previous investigation showed that the R^2^ for predicted and observed *in vivo* log CL_int_ for human hepatocyte CL_int,u_-based predictions was 0.3-0.4 [21, 25]. Stringer et al. [33] found a R^2^ that appeared to be lower than that, both for hepatocyte and (in particular) microsome CL_int,u_- based predictions.

Using a test set of 73 drugs, we found that the R^2^ between predicted and observed *in vivo* log CL_int_ for human microsome CL_int,u_-based predictions was 0.18 (Figure 2) [30]. In these microsomal experiments, drugs such as naloxone, lidocaine, glipizide and ondansetron with moderate to moderately high *in vivo* CL_int_ (*in vivo* CL_int_ corresponds to *in vitro* CL_int,u_) were found to have *in vitro* CL_int,u_<LOQ. The maximum prediction error was 350-fold (overprediction). *In vivo* CL_int_ threshold for proposed low CL_int_ ranged between 15 and 1,900 mL/min. The average and maximum intra-laboratory variability were 6.7- and 58-fold [10, 30, 34], respectively. Every other compound had individual microsome CL_int,u_-measurements in more than one class (at least two measurements were done for each compound), and 10 % of them had individual CL_int,u_- measurements spanning over 2-3 classes (one compound, alfentanil, had low and very high CL_int,u_ on different occasions). Thus, the reproducibility of the microsome assay is limited. For every other compound *in vitro* and *in vivo* CL_int_-classes differed [10, 30]. Ca 30 % of compounds with low measured microsomal CL_int,u_ had low *in vivo* CL_int_, and 67 % of compounds with low *in vivo* CL_int_ had low *in vitro* CL_int,u_. which shows an overprediction trend.

By adding human microsome CL_int,u_ for 7 OATP-substrate drugs (with available *in vivo* CL_int_ data) from [35] to the set of 73 compounds, the R^2^ decreased to 0.11 [30]. The *in vivo* CL_int_ for OATP-substrates was on average more than 10-fold higher than predicted from microsome CL_int,u_ [35].

Due to LOQ-limitations, a substantial fraction of compounds is normally excluded from such correlations and evaluations [25, 33, 41]. Stringer et al. [33] found that none of their selected test compounds with *in vivo* CL_int_ <70 mL/min had quantifiable microsomal *in vitro* CL_int,u_. and that 8, 31 and 75 % of compounds with *in vivo* CL_int_ 70-700, 700-7,000 and >7,000 mL/min had quantifiable microsomal *in vitro* CL_int,u_, respectively [33]. Thus, there is an approximately 50 % chance that a compound with *in vitro* microsomal CL_int_<LOQ has high to very high *in vivo* CL_int_.

The estimated median *in vivo* CL_int_ and our proposed limit for low CL_int_ for drugs are 850 and 150 mL/min, respectively. The corresponding limit for low CL_int_ that was set by Sohlenius-Sternbeck and Terelius (ca 2,000 mL/min) [10] is about 3- and 13-fold higher than these, respectively. Corresponding thresholds for very high CL_int_ are ca 20,000 and 50,000 mL/min (2.5- fold lower *in vitro* [10]), respectively. Thus, limits proposed for microsome CL_int,u_ are relatively high at low CL_int_ and low at high CL_int_.

For metabolically unstable compounds, *in vitro* t_½_ could be <10-20 min (>100 µL/min/mg), which implies higher uncertainty of measured and predicted CL_int_.

Among top 5 % CL_H_-drugs, less than 1/3 belong to those with highest CL_int_, showing that also f_u_ (and CL_int_ • f_u_) are important parameters for metabolic stability *in vivo*. For those with the highest estimated CL_H_, none belong to the very high CL_int_-class. Peak CL_H_ is found at CL_int_=24,000 mL/min, which is about 1/10 of the maximum estimated *in vivo* CL_int_.

The mean fraction excreted renally (f_e_) at *in vivo* CL_int_ of 150 (our proposed threshold for low CL_int_), 850 (median CL_int_ for drugs), 2,000 (approximate threshold for low CL_int_ according to [10]), 5,000 (approximate threshold for very high CL_int_ according to [10]) and 50,000 (our proposed threshold for very high CL_int_) mL/min are 54, 27, 19 (range 0-85 %), 13 (range 0-70 %) and 2 %, respectively.

Interlaboratory variability for unbound microsomal fraction (f_u,mic_), which is used to convert CL_int_ to CL_int,u_, is also a source of uncertainty for CL_int_-predictions. For example, ratios between highest and lowest reported f_u,mic_-estimates for amitriptyline and imipramine are 5- to 6-fold. *In silico* prediction models for f_u,mic_ reach R^2^ of ca 0.6, with 1.5 average-fold error and some cases with 10- to 100-fold prediction errors [31]. This sets the upper accuracy limit for CL_int,u_- predictions.

#### Blood-brain barrier permeability and brain uptake

An evaluation of BBB permeability and uptake demonstrated that this barrier is highly permeable, sufficient to absorb compounds with MW>1000 g/mol, log D<-3.5 and polar surface area >270 Å^2^ well [37]. The *in vitro* BBB P_app_ is higher than in Caco-2 cells [38], on average 11-fold and maximally 34-fold higher.

Brain uptake index-data from rats *in vivo* showed rapid and extensive brain uptake of high permeability compounds during a very short experimental period and some absorption of low permeability substances [37]. For example, antipyrine, caffeine, nicotine and propranolol were absorbed to ∼70-100% within 5-15 s, and the uptake of hydrocortisone and sucrose was 1.4 % [37].

The BBB passage times for different molecules (from the low permeability compound sucrose to the highly permeable propranolol) in capillary endothelial BBB cells were approximated to ∼0.1 to ∼4 s [37]. Corresponding *in vivo* estimates obtained in rats *in vivo* were ∼0.3 s to ∼12 min (0.2 s, 7 s and 50 min for caffeine, morphine, and inulin (MW∼5000 g/mol), respectively). In humans *in vivo*, many highly permeable substances, including anaesthetics and nicotine, have a very rapid on-set of CNS effects (in the order of seconds) following injection or inhalation. Morphine, with moderate passive BBB permeability, significant BBB efflux (by MDR-1; ratio between unbound brain and plasma concentrations (K_p,uu,brain_) in rats = 0.15) and low binding capacity to brain tissue has an on-set time of 5-10 min. The transport through the gut-wall, rather than the passage across the BBB, appears to be rate-limiting. Remifentanil has a blood-brain equilibration t_½_ of 1 min and a rapid onset of action (general anesthesia) in humans.

K_p,uu,brain_-estimates obtained in rats and humans differ. A 0.01 correlation between the two was found [42]. Among reasons to the poor relationship include the relatively high expression of mdr-1 and low expression of bcrp in rats (compared to human MDR-1 and BCRP), 15 % difference in MDR-1/mdr-1 homology between humans and rats, and overall higher plasma f_u_ in in rats [42]. One compound with high K_p,uu,brain_ in man (2.8) is effluxed and predicted to have high passive permeability, but has no apparent efflux at the rat BBB. A MDR-1 substrate with high K_p,uu,brain_ in rat (2.4) has very low K_p,uu,brain_ in man (0.15) and is predicted to have high passive permeability.

A high log BB is (according to the BBB+BBB- concept) indicative of high BBB permeability and CNS-activity potential. There are, however, CNS-active compounds with very low log BB in rats (for example, log BB<2 (BB<0.01)) [37].

### *In silico* and Physicochemical Properties to *in vivo* Predictions

#### Log D to oral bioavailability (F) and half-life (t_½_)

Figures 4 and 5 show the correlations between measured log D and F (n=212; R^2^=0.23; corresponding R^2^ for predicted log D *vs* F=0.11) and log t_½_ (n=423; R^2^=0.065; corresponding R^2^ for predicted log D *vs* log t_½_=0.035), respectively. Note that F- and/or log D-values for many highly lipophilic compounds are not available.

**Figure 4.**
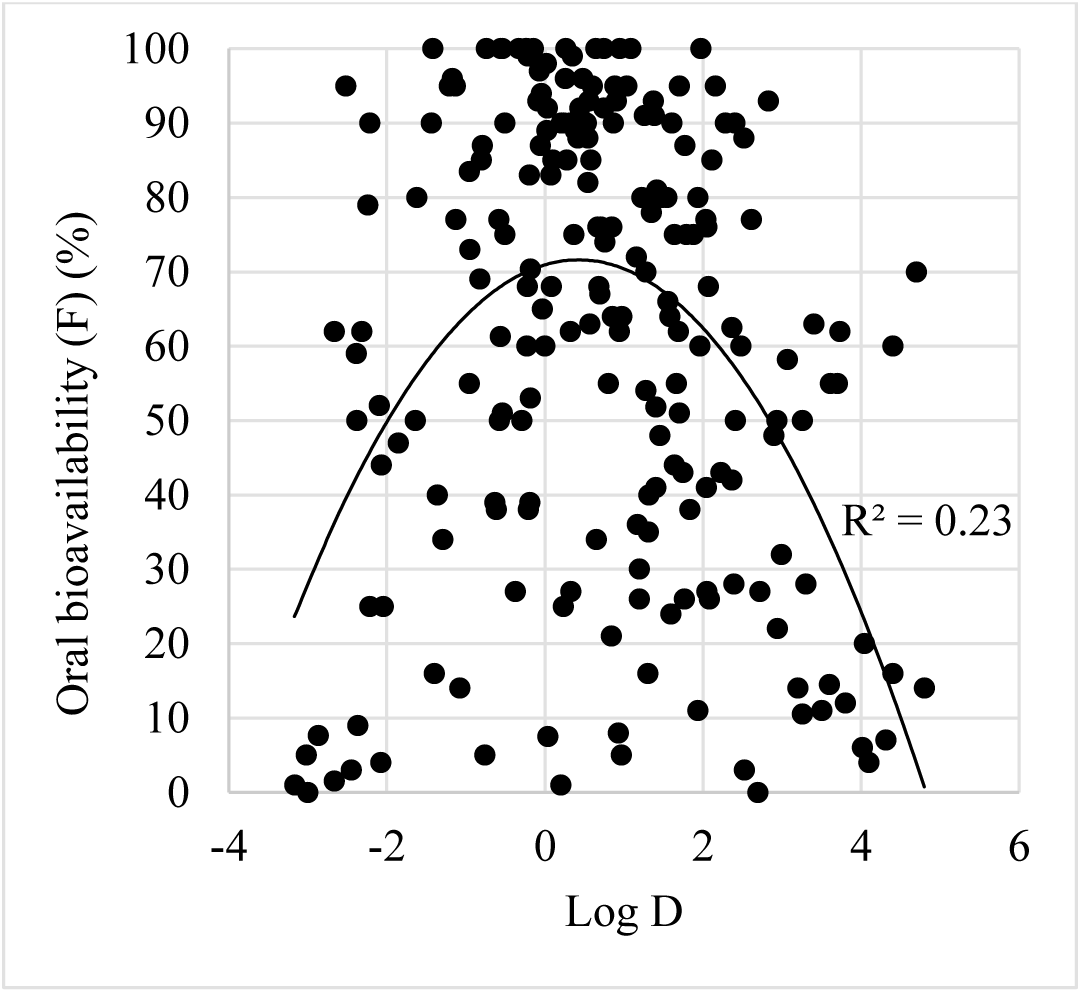
The correlation between measured log D and oral bioavailability (F) for 212 compounds with log D ranging from −3.2 to 4.8 and F ranging from ca 0 to ca 100 %. For predicted log D *vs* F the R^2^ was 0.11.

**Figure 5.**
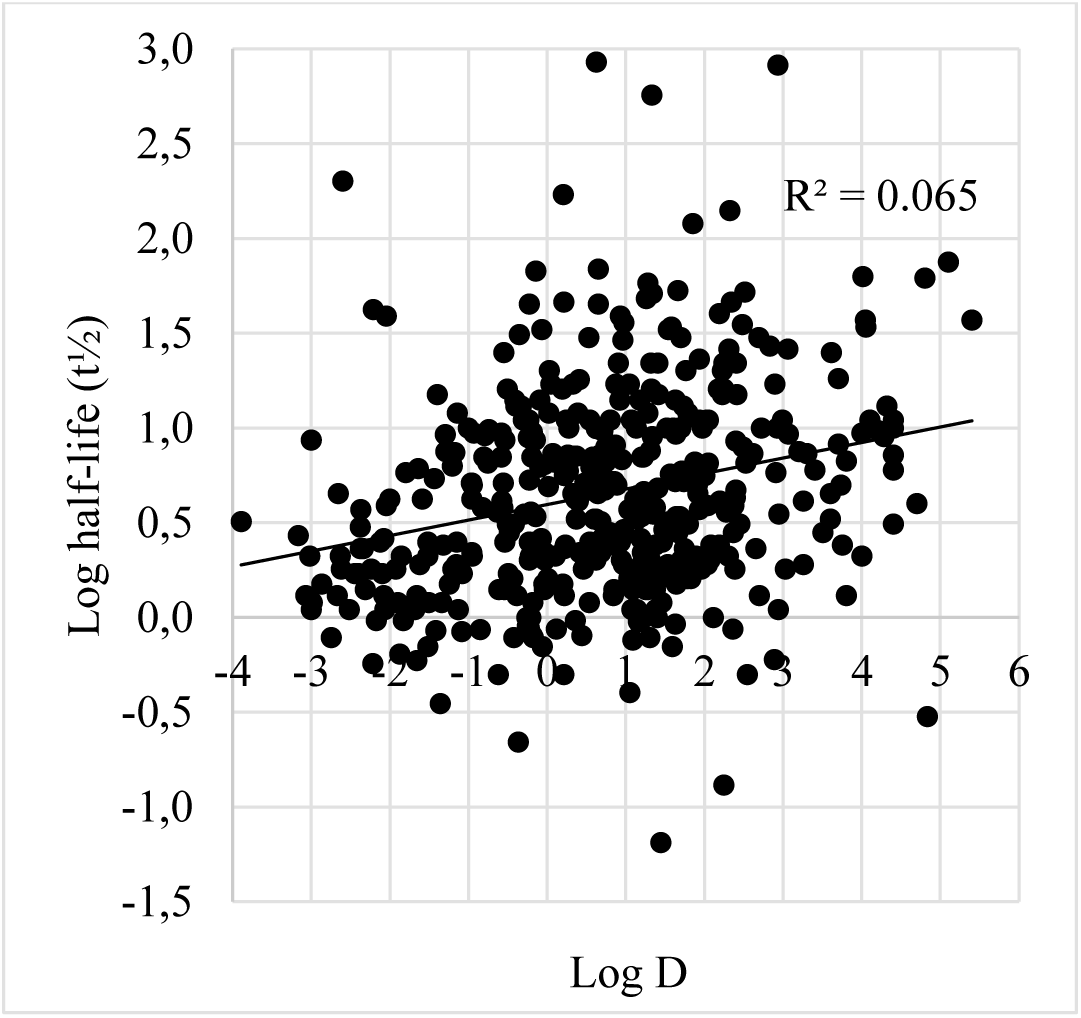
The correlation between measured log D and log half-life (t_½_) for 423 compounds with log D ranging from −3.9 to 5.4 and t_½_ ranging from 4 min to 35 days. For predicted log D *vs* log t½ the R^2^ was 0.035.

According to the fitted relationship between log D and F, the optimal log D for achieving the highest possible F is ca 0.5. At log D 0-1 there are, however, compounds with very low F (<10 %), and >70 % F has been reached for compounds with log D from −2.5 to 4.7. At log D 0-1 estimated t_½_ ranges from 25 min to 35 days.

#### Rule of 5 (Ro5) and fraction absorbed (f_abs_)

Among a collection of 129 Ro5 violent compounds (2 or more of the rule criteria violated) we found 59 with available f_abs_-values. The mean f_abs_ for these was 42 % (compared to ca 80 % for compounds with Ro5 acceptance) [36]. Only 37 % of these had poor f_abs_ (<10 %) and 15 % had a f_abs_ exceeding 90 %. One compound with 3-4 violations of the rule (tenapanor) is absorbed to 22 % and one compound with log P of 8.2 (venetoclax) has a f_abs_ of >65 %. Out of 77 selected compounds in the study that do not violate more than 1 Ro5 criteria 3 (4 %) were found to be false negatives (having f_abs_<10 %).

Suenderhauf et al. built *in silico* models for the prediction (rather fitting, since test set was apparently missing) of f_abs_, and reached a R^2^ of 0.6 and a RMSE of 26 % (n=458) [40]. The intercept for observed *vs* predicted f_abs_ deviated, however, markedly from zero, and was as high as ca 50 % f_abs_. Ca 20 % of compounds with f_abs_<10 % were predicted to have f_abs_<10 % and ca 40 % of these compounds were predicted to have f_abs_>50 %. 10 compounds with predicted 90-98% f_abs_ are poorly absorbed *in vivo* (including 2 with f_abs_<5 %), showing risk of selecting compounds with unacceptable uptake properties.

With class models (only f_abs_<30 % or >80 %), the Receiver Operating Characteristics (ROC) averaged ca 0.7 [40]. Numerical models by other groups reached R^2^ (no forward-looking predictions) and RMSE of 0.7-0.9 and 14-35 %, respectively.

With ADMET Predictor (where f_abs_ is predicted via P_eff_, and many of the test set compounds probably were present in the training set), a R^2^ of 0.1 and an intercept of roughly 40 % f_abs_ (overprediction trend at low f_abs_) were reached (n=several hundred; data on file). This R^2^ is of same size as found for *in silico* predictions of F with ADMET Predictor (R^2^ ca 0.15 for compounds with complete absorption, and likely not more than half of that if including incompletely absorbed compounds) with this software [19].

### *In silico* to *in vitro* to *in vivo* Predictions and Thresholds

#### Solubility to in vivo dissolution (f_diss_)

The Q^2^ for *in silico vs in vitro* log S (including ADMET Predictor) and R^2^ for log S *vs in vivo* f_diss_ were ca 0.4 (range 0.13-0.8) and 0.13, respectively (Figure 2). The corresponding *in silico-in vitro* Q^2^ - *in vitro-in vivo* R^2^ product is 0.05. The Q^2^ • R^2^ applies within the moderately low to very high S range. Sufficient information for determination of Q^2^ • R^2^ for compounds with low S (<1- 10 µM) and incomplete f_diss_ is lacking.

An overprediction trend at low S (ca 95 % of low S compounds were *in silico* predicted to have moderate to very high *in vitro* S with ADMET Predictor [10]), lack of many S-values due to LOQ-limitation, and a relatively high set *in vitro* threshold for poor/low S (apparently, roughly at least 15-fold higher than found for *in vivo* f_diss_; 10 *vs* 0.5-1 µM) causes uncertainty for prediction of *in vivo* f_diss_ from S.

For compounds with predicted very high S, ca 67 % have very high measured S, and ca 8 % have poor S (<10 µM) (according to ADMET Predictor results by [10]), and at the same time, 10 µM often consistent with complete *in vivo* f_diss_.

#### Permeability to fraction absorbed (f_abs_)

The overall Q^2^ for *in silico vs in vitro* log permeability and R^2^ for log P_app_ *vs in vivo* f_diss_ were both ca 0.6 (range ca 0.2-0.8) (Figure 2). The corresponding *in silico-in vitro* Q^2^ - *in vitro-in vivo* R^2^ product is 0.36. The Q^2^ • R^2^ applies within the moderately high to very high permeability and f_abs_ range (>80-90 % f_abs_). At low and moderate permeability and f_abs_ (<80-90 %), close to zero Q^2^ (and Q^2^ • R^2^) was found. *Note: It is possible that compounds without in vitro P_app_ due to low experimental recovery were excluded*.

Corresponding *in silico vs in vitro* Q^2^ and Q^2^ • R^2^ for ADMET Predictor, where P_eff_ is an intermediate between P_app_ and f_abs_, was lower, ca 0.45 and ca 0.25 (roughly estimated), respectively [10]. With a similar approach (MDCK P_app_ to *in vivo* f_abs_ via P_eff_), 0.43 and ca 0.2, respectively [31].

Using ADMET Predictor, ca 70 % of compounds with predicted (proposed) low P_eff_ (<1 • 10^-4^ cm/s; corresponds to ca <90 % f_abs_ [32]) had measured P_app_<2 • 10^-6^ cm/s (which corresponds to ca <65 % according to [23]). P_eff_- and P_app_-thresholds for low *in vivo* permeability are <0.1-0.3 • 10^-4^ cm/s (<ca 30-60 % f_abs_ [32] and <0.1 • 10^-6^ cm/s (f_abs_<ca 30-40 % [23]) rather than the 3- to 10- and 20-fold higher limits proposed (by [10]), respectively. Ca 13 % of compounds with predicted P_eff_<1 • 10^-4^ cm/s had very high measured P_app_ (>10 • 10^-6^ cm/s), and ca 67 % of compounds with predicted very high P_eff_ also had very high measured P_app_. None of the compounds with predicted very high P_eff_ (>3 • 10^-4^ cm/s) had measured P_app_ <2 • 10^-6^ cm/s).

#### Intrinsic clearance (CL_int_)

The Q^2^ for *in silico vs in vitro* log CL_int_ using ADMET Predictor and R^2^ for microsomal *in vitro* CL_int_ *vs in vivo* log CL_int_ were 0.32 (mean of 0.50 and 0.14) and 0.15 (based on R^2^ values with and without OATP-substrates from different studies, excluding compounds with CL_int,u_<LOQ), respectively (Figure 2). *In silico-in vitro* Q^2^ • *in vitro-in vivo* R^2^ = 0.05 (for compounds with microsomal CL_int_>LOQ).

Metabolic stability *in vivo* (e.g. CL_H_) is also determined by f_u_. Based on prediction accuracies for both CL_int_ (0.15 for *in vitro* to *in vivo*; 0.05 for *in silico* to *in vitro* to *in vivo*) and f_u_ (0.45 for *in silico* to *in vitro*), CL_H_ is also likely to be poorly predicted.

The corresponding *in vivo* CL_int_ for the proposed low microsomal CL_int_ (<15 µL/min/mg) is approximately 3 times higher than for the typical marketed drug (2,000 *vs* 850 mL/min), and considered moderately low to moderate. What was considered very high and undesirable for microsome CL_int_ is ca 2.5-fold lower than the proposed threshold for the *in vivo* situation (ca 20,000 *vs* 50,000 mL/min).

Ca 18 % of compounds with *in silico* predicted proposed low CL_int_ had proposed very high and undesirable measured *in vitro* CL_int_, and the percentage in the opposite direction was ca 26. In contrast, there was an underprediction trend at low CL_int_ for *in vitro* to *in vivo* as only ca 30 % of compounds with low microsomal CL_int_ had low *in vivo* CL_int_.

## DISCUSSION

In this investigation it is apparent that commonly used early discovery ADME/PK numerical and classification models (including the ADMET Predictor software) and rules of thumb have poor validity (poor accuracy, limited range, systematic errors, very low/no clinical relevance) and inadequate thresholds for poor/acceptable values.

Poor reliability of *in silico* to *in vitro* to *in vivo* and *in silico* to *in vivo* predictions was shown by low accuracy (Q^2^ • R^2^) for log S to f_diss_ (0.05), log permeability to f_abs_ (0.36; 0.25 for the ADMET Predictor approach that includes the intermediate P_eff_-step), log CL_int,u_ to log CL_int_ (0.05), log CL_H_ (no number estimated, but apparently, very low), log D *vs* F (0.11) and log t_½_ (0.035). At low to moderate, often decision-critical, levels, Q^2^ • R^2^-estimates were ∼0.

The low accuracies and limited ranges for CL_int_ and f_abs_, and skewness and limited ranges for S- and f_u_-models, indicate that CL_H_, CL, F and t_½_ must also be poorly predicted. *Note: The assumption that Q^2^ • R^2^ correctly reflects the true in silico to in vivo predictive power might not be accurate. True values might be somewhat lower or higher*.

The average Q^2^ for *in silico* to *in vitro* log S, log P_app_ and log CL_int_ was estimated to 0.44, which was higher than the corresponding mean R^2^ for *in vitro* to *in vivo* (0.26). Thus, most of the predictive accuracy from *in silico* to *in vivo* was lost in the translation between *in vitro* and *in vivo*. Furthermore, application/prediction range is also lost with *in vitro* data. For example, low S and f_u_ and low to moderate (sometimes up to high) microsomal CL_int_ is often not quantifiable.

Thus, unless predicted S, permeability and f_u_ are high and CL_int_ is moderate one is more or less lost in the translation and predictions and will jeopardize compound selection and optimization. Apparently, none of 28 modern small drugs (based on drugs and data from 2021) falls into this character [18]. Eighteen % of them did not meet any of these criteria, and nearly half of them only reached maximum one criterium (Figure 6) [18]. The typical (based on median observed or predicted values for small drugs marketed in 2021 [18]) modern small drugs has a f_diss_ of 95 % (60 % with *in vivo* dissolution limitation), f_abs_ of 70 % (83 % with *in vivo* f_abs_<90 %), f_u_ of 4 % (52 % with f_u_<5 %) and CL_int_ of 2,500 mL/min (45 % with very high *in vitro* CL_int,u_ according to previously established limit) [18]. It is therefore highly probable that at least one of *in silico* predictions of S, P_app_, CL_int,u_ and f_u_ for modern small drug candidates will be very poor (uncertain, overpredicted and/or underpredicted). This will likely lead to erroneous interpretations and decisions and shows the need for other better tools. The findings and/or predictions that significant renal excretion, gut-wall extraction and/or bile excretion are involved for 30-55 % of the new drugs show additional obstacles and needs [18].

**Figure 6.**
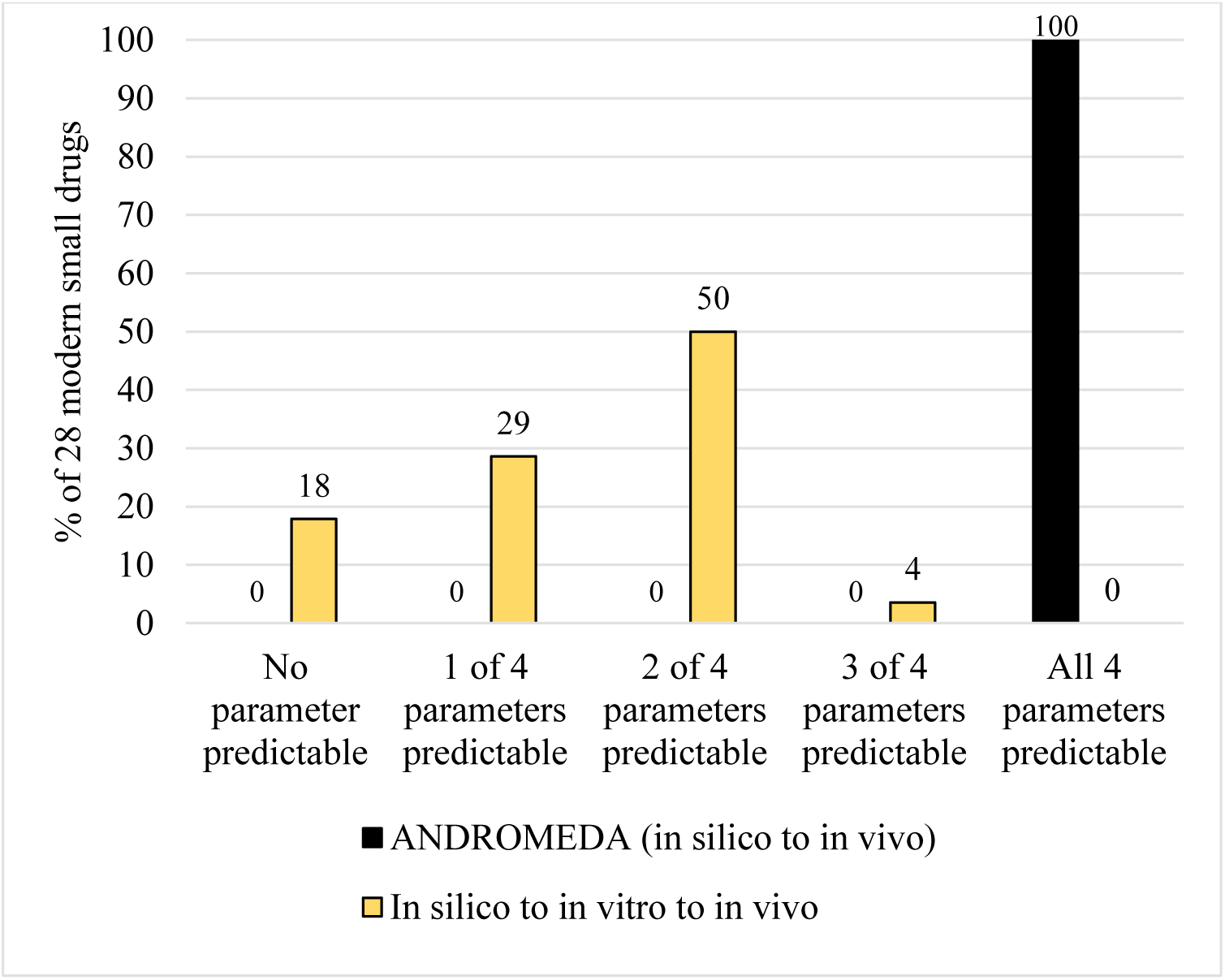
Estimated coverage (prediction ranges for solubility/dissolution, permeability/absorption, metabolic instability and unbound fraction in plasma) of *in silico* to *in vitro* to *in vivo* (for example, ADMET Predictor) and ANDROMEDA (*in silico* to *in vivo*) prediction methodologies for 28 modern small drugs (marketed in 2021).

Due to poor to moderately high R^2^ and Q^2^, there were also quite high percentages of incorrect predicted class, 43 to 63 % incorrect for *in silico* to *in vitro* log S (in particular for low S compounds), f_abs_ (in particular for high P_app_ compounds) and CL_int_ (Figure 1).

Thresholds for poor S (low f_diss_), poor permeability (low f_abs_) and undesirably high CL_int_ are lower (at least ca 15-fold for S), lower (ca 10-fold for P_eff_ and ca 20-fold for P_app_) and higher (ca 2.5-fold for CL_int_) according to our *in vivo* data based proposals. An adjustment to these is assumed to lead to better decision-making (stop/go).

Higher accuracy was demonstrated for *in silico* predictions of *in vivo* f_abs_ (0.1-0.6) and *in vitro* f_u_ (0.4-0.5). However, it could not be ruled out that the presence of training set compounds in test sets had exaggerated the predictive power, and both models were shown to not be applicable for compounds with lower estimates. Lack of correlation between predicted and observed values and marked skewness at low permeability and f_abs_ (large overprediction trend at low f_abs_) limits the applicability of numeric *in silico* f_abs_-models, whereas lack of highly bound (f_u_<1 %; including many compounds with f_u_<LOQ) in evaluations and marked skewness (large overprediction trend at f_u_<25 % and underprediction trend at high f_u_) limits the applicability *in silico* the investigated f_u_-prediction models in [17]. Since many modern small drugs have f_u_ of maximally 2 % (for ca 30 % of small drugs; [18]) and/or limited permeability and f_abs_ (for nearly every other small drug [18]), this is believed to have a major negative impact on the applicability of these models.

Limited range and skewness were also evident for S and CL_int,u_ at the lower values. The portion of non-quantifiable compounds was particularly high for CL_int,u_ (up to every other compound). A belief that 10 µM generally corresponds to solubility/dissolution problems *in vivo* and that an *in vitro* CL_int,u_ of 15 µL/min/mg is low and reflects good metabolic stability leads to an underestimation of dissolution potential and overestimation of metabolic stability *in vivo*. This might cause incorrect decision-making, compound selection and optimization strategies.

The clinical relevance and validity are also poor for aqueous S and microsomal CL_int,u_. The R^2^s for log aqueous S *vs* S in simulated (FaSSIF) and real (HIF) intestinal fluids and *in vivo* f_diss_ are estimated to 0.31, 0.15 and 0.13, respectively (data on file). The low R^2^s for log S reflect the differences between media regarding content, dynamics and pHs. There are also marked differences between incubated microsomes and hepatocytes in the liver *in vivo*. Microsomes lack an outer cell membrane and its transporter proteins and lack some binding components. They show reduced enzymatic activity (especially for conjugation) and there is considerably smaller ratio between cells/cell components (microsomes *in vitro* and hepatocytes *in vivo*) and fluid (incubation fluid *in vitro* and blood *in vivo*). There is also uncertainty how to compensate for binding capacity (f_u,mic_ is used for conversion of microsome CL_int_ to CL_int,u_, but its clinical relevance is questioned) and detaching rates from binding sites. The Caco-2 cell model appears closer to the *in vivo* situation, which is also indicated by a relatively high *in vitro-in vivo* correlation.

Due to the importance of f_u_ for CL_H_ (a low f_u_ compensates for a high CL_int_ *in vivo*) and the observation that it is generally not the compounds with highest CL_int_ that have the highest CL_H_, it is our proposal that CL_int_ • f_u_ (or CL_H_) should replace CL_int_ as a measurement and classification of metabolic instability and that *in silico* models for f_u_-predictions include compounds with very low values.

The importance of f_u_, steady-state volume of distribution (V_ss_) and excretion for overall CL and t_½_ is apparent. Some drugs with very good metabolic stability (very low *in vivo* CL_int_; probably <LOQ for microsomes) have short t_½_ (<2 h) - captopril, carboplatin, inogatran, adefovir, amoxicillin, aztreonam, cefuroxime, melagatran and metformin, to name a few. On the contrary, some drugs with very high *in vivo* CL_int_ have long t_½_ (days to weeks). These include clomipramine, itraconazole and amiodarone.

Other drugs worth mentioning are atenolol (low *in vivo* CL_int_; zero, low and moderate microsomal CL_int,u_ reported [30]), diclofenac (high *in vivo* CL_int_; very high microsomal CL_int,u_; low to high hepatocyte CL_int,u_ (57-fold difference between lowest and highest reported values) [30, 43, 44]), gemfibrozil (moderate *in vivo* CL_int_; moderate microsomal CL_int,u_; low to high hepatocyte CL_int_ (138-fold difference between lowest and highest reported values) [30, 43, 44]), and telmisartan (very high *in vivo* CL_int_; zero and low microsomal CL_int,u_ (>200-fold underprediction) [35]). *In vivo* in man, telmisartan is a very high CL_int_, very low f_u_ drug, but it appears likely that *in silico* models would predict limited microsomal CL_int,u_ and much higher f_u_ than estimated *in vitro* (using ADMET Predictor or models by Watanabe et al. and Ingle et al.).

Earlier proposed ideal log D for ADME/PK (log D of 0-3 or 2-3) got support by the peak F found at log D=0.5 (Figure 4). However, the correlation between log D and F is weak, many compounds within the range have poor F, and outside the proposed optimum zone there are several compounds with good F. An even weaker correlation was found for log D and log t_½_ (Figure 5). Thus, this rule of thumb is not particularly reliable for these two secondary ADME/PK parameters.

The evaluation of Ro5 for classification of poor and adequate f_abs_ (<10 or >10 % f_abs_) showed that this rule is skewed and produces a large portion of false negatives and a minor portion of false positives. 63 % of drugs classified to have poor absorption showed moderate to complete f_abs_ *in vivo*. Thus, the application of log D-, MW- and HB-dependent Ro5 implies a substantial risk to opt out compounds with sufficiently good oral absorption potential *in vivo* in man.

The BBB+/BBB- concept (based on rodent log BB-data and thresholds) does not seem valid and useful either. This is because the BBB appears highly permeable, at least for small drugs, log BB is highly dependent on the binding to brain and blood components, species differences in brain uptake capacity and efflux exist, and there are many CNS-active compounds with very low log BB (belonging to proposed BBB- class) [37]. K_p,uu,brain_ is a more relevant parameter for assessing and describing uptake capacity across the BBB [42]. However, an investigation showed no correlation between rat and human K_p,uu,brain_-values [42]. Whether this was mainly due to experimental factors or differences in efflux transporter expression and activities is unknown. The BBB permeability and efflux (mainly MDR-1 and BCRP) and brain binding reference system developed by us, included in the ANDROMEDA prediction software, is an alternative [45].

ANDROMEDA is an alternative to the *in silico* to *in vitro* to *in vivo* approach. This software is mainly based on models built and trained on human clinical data and predicts directly from chemical structure to *in vivo* in man (without the need and use of *in vitro* data for S, P_app_ and CL_int,u_). Advantages compared to *in vitro* data-based *in silico* models (including ADMET Predictor) include wider application domain (high resolution at low values), higher accuracy, more parameters (including renal and biliary CL and gut-wall extraction), low/minimal skewness, high clinical relevance, lower variability/uncertainty of data used for model training and clinically more relevant thresholds. Q^2^-values (corresponding Q^2^ • R^2^- values for *in silico-in vitro-in vivo* within parentheses) for f_abs_, f_diss_, log CL_int_ and f_u_ are 0.7-0.8 (0.36), 0.55 (0.05), 0.55 (0.05) and 0.5-0.55 (0.4-0.5), respectively. ADMET Predictor reached a Q^2^ of 0.15 for F for compounds without absorption limitations (assumed to be <0.15 including compounds with incomplete absorption). The mean Q^2^ for F with ANDROMEDA, reached in 4 different validation and benchmarking studies, was 0.49 [19]. The overall mean values for these 5 parameters are 0.57 (0.2) (3 times higher for ANDROMEDA). Corresponding prediction and application ranges with ANDROMEDA are 0-100 % (ca 0-100 %), 1-100 % (higher than 1 %, but unknown, to 100 %), 1-200,000 mL/min (roughly 50-5,000 mL/min to ca 80,000 mL/min), 0.02-100 % (ca 25-100 %) and 0-100 %, respectively. Other advantages with ANDROMEDA include additional essential parameters, such as renal and biliary CL, gut-wall extraction ratio, blood-plasma concentration ratio, V_ss_, t_½_, CYPID and efflux transporter ID, and unique 70 % confidence intervals predicted for each compound and parameter estimate. The latter implies that measures of confidence are produced. ANDROMEDA was very successful in predicting the human clinical ADME/PK for the small drugs released on the market in 2021 and a challenging set of compounds proposed for benchmarking and validation of prediction methods (Figure 6) [18]. It predicted f_diss_, f_abs_, CL_int_ and f_u_ for all these drugs, and the average and maximum individual prediction error for all investigated parameters were 2.6- and 16-fold, respectively [18].

With traditional models and thresholds there is an imminent risk to incorrectly opt out compounds and select compounds that (later) will show poor ADME/PK-characteristics. That is a matter of costs, time, uncertainties and risks. These risks are considerably lower with ANDROMEDA.

## CONCLUSION

The validity of investigated methodologies (including ADMET Predictor) and thresholds were overall very low, and much of the predictive power was lost by including and predicting via *in vitro* data. Unless both predicted S, permeability and f_u_ are high and CL_int_ is moderate (an overall criterium not met for any of the investigated modern small drugs) one is more or less lost in the translation and predictions, and this will likely lead to erroneous interpretations and decisions and jeopardizing of compound selection and optimization. It is surprising that these findings have not been shown before. This clearly shows the need for better and thoroughly validated prediction models and software. Marked improvements in accuracy, range, balance and clinical relevance were achieved with ANDROMEDA, which predicts more than 30 human clinical ADME/PK parameters directly from chemical structures and has undergone extensive validation and benchmarking. ANDROMEDA successfully predicted the human clinical ADME/PK for drugs marketed in 2021, and for which the *in silico* to *in vitro* to *in vivo* approach seems out of reach.

